# Cav1.3-C-terminus Restores Electrical and Contractile Functions in hiPSC-derived Cardiomyocytes Model of Heart Failure

**DOI:** 10.64898/2026.10.27.762268

**Authors:** Yvonne Sleiman, Jean-Baptiste Reisqs, Mohamed Boutjdir

## Abstract

**Objective:** Heart failure (HF) is a major global health concern characterized by impaired excitation-contraction coupling and extensive structural remodeling of the myocardium, often modeled using a potent vasoactive and neurohumoral mediator, endothelin-1 (ET-1). This study investigates whether AAV6-mediated delivery of the Cav1.3-C-terminus, acting as an auto-trans-regulator of gene transcription, can restore electrophysiological and contractile functions in an ET-1-induced HF model using hiPSC-CMs.

**Methods:** hiPSC-CM monolayers were treated with 10 nM ET-1 for 48 hours to induce HF-like phenotype and then co-treated with an AAV6-Cav1.3-C-terminus. We subsequently evaluated gene expression via RT-qPCR, single-cell electrophysiological properties via patch-clamp, and contractile function using custom video analysis software.

**Results:** ET-1 treatment induced a HF-associated remodeling phenotype, characterized by significant upregulation of hypertrophic (*NPPA*, *NPPB*, and *ACTA1*) and extracellular matrix-associated (*COL1A1*) genes, along with increased cell membrane capacitance. This structural remodeling was accompanied by marked electrophysiological alterations, including downregulation of *KCNH2*, *CACNA1D*, *CACNA1C*, and *SCN5A*, reduced potassium, calcium, and sodium current densities, and prolonged action potential duration. ET-1 impaired both the spontaneous beating rate and contraction amplitude of the monolayers. Co-treatment with the Cav1.3-C-terminus successfully restored *CACNA1D* and *KCNH2* gene expression to baseline levels, accompanied by normalized L-type calcium and potassium current densities, normalized APD prolongation, and completely restored the chronotropic and inotropic functions to control levels.

**Conclusions:** ET-1-treated hiPSC-CMs model effectively recapitulates the electrophysiological and contractile dysregulations of HF. Gene therapy by Cav1.3-C-terminus provides robust functional rescue myocardial contractility, highlighting its potential as a novel therapeutic strategy for HF.

**Graphical Abstract:** 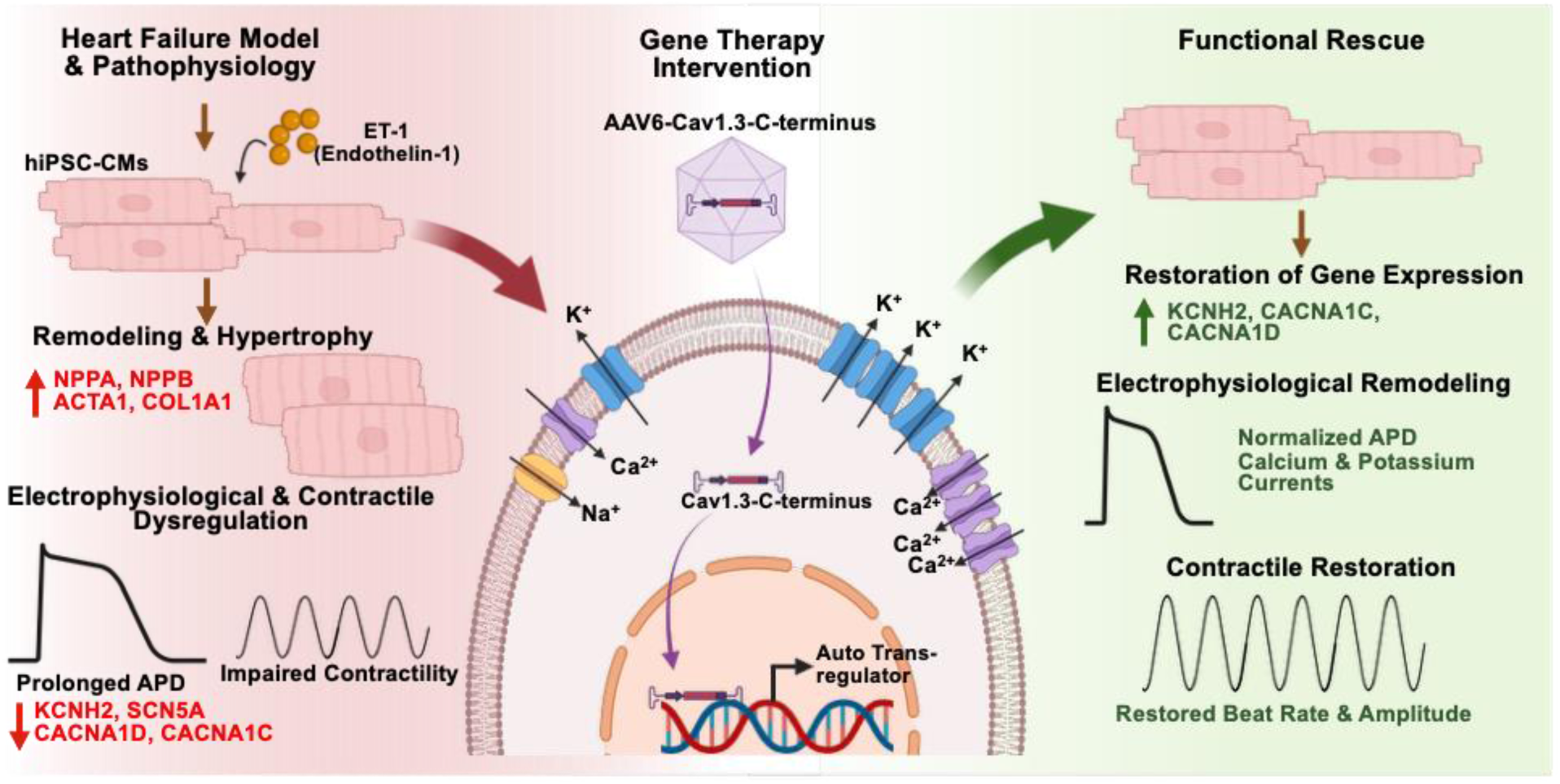

**Significance statement:** Heart failure (HF) involves structural, electrical and contractile dysfunctions with limited curative options. This study highlights a novel gene therapy utilizing the auto-trans-regulatory properties of the Cav1.3 C-terminus. In an endothelin-1-induced HF model using human induced pluripotent stem cells-derived cardiomyocytes (hiPSC-CMs), AAV6-mediated delivery of the Cav1.3-C-terminus successfully reversed pathological remodeling. By normalizing action potential duration, critical ion channel expression (CACNA1D, CACNA1C KCNH2), and rescuing mechanical contractility, this gene therapy approach offers a highly promising, targeted therapeutic strategy for tackling the complex pathophysiology of heart failure.

## Introduction

Heart failure (HF) remains a major global health concern and medical challenge, affecting approximately 64 million people worldwide, with its prevalence in the United States projected to reach 8.5 million by 2030.^1,2^ Often resulting from ischemic coronary disease or microvascular dysfunction, cardiac valve defect, diabetes, or inherited cardiomyopathy, HF is characterized by the heart’s inability to pump blood efficiently enough to meet the body’s metabolic demands, leading to symptoms such as fatigue, shortness of breath, and fluid retention in the lungs.^3^ During myocardial ischemia, reduced oxygen supply compromises cardiomyocyte function and, if sustained, leads to cell death and impaired myocardial contractility. In response, the surviving myocardium undergoes compensatory hypertrophic remodeling to preserve cardiac function. However, prolonged activation of this initially adaptive response can lead to maladaptive cardiac remodeling and progressive deterioration of cardiac function.^4^ This remodeling manifests as sarcomere disorganization, fibrosis, metabolic imbalances, and critically a profound dysregulation of intracellular calcium homeostasis.^5,6^ In the failing heart, excitation-contraction coupling is severely compromised, characterized by the depletion of sarcoplasmic reticulum (SR) calcium stores. This depletion is driven by the downregulation of the SERCA pump and the pathological hyperphosphorylation of ryanodine receptor 2 (RyR2), which creates a continuous diastolic calcium leak.^7^ Consequently, the decreased of systolic calcium amplitude directly impaired myocardial contractile force, while the diastolic calcium overload impairs the relaxation.

Pathological hypertrophy is closely linked to the overproduction of neurohumoral mediators, including endothelin-1 (ET-1).^8,9^ This potent vasoactive and cardioactive peptide activates G protein-coupled receptor (GPCR) on the cardiomyocyte membrane, acting as a key initiator of maladaptive hypertrophic remodeling.^10^ ET-1 also triggers the expression of stress genes, causing the accumulation and release of natriuretic peptides (ANP and BNP) that attempt to counteract the pathological activation of renin-angiotensin-aldosterone and sympathetic nervous systems.^11,12^ Due to its ability to faithfully recapitulate these molecular pathways, ET-1 has been used to model HF *in vitro.*^13,14^

Although small- and large-animal models have provided valuable insights into the mechanisms underlying HF, their translational relevance is limited by species-specific differences in cardiac physiology, electrophysiology, and disease progression. Consequently, many mechanistic studies rely on *in vitro* models, including primary cardiomyocyte cultures and established cell lines, which, despite their experimental advantages, often fail to fully recapitulate the structural, functional, and molecular complexity of human cardiac disease.^15^ The integration of human induced pluripotent stem cell-derived cardiomyocytes (hiPSC-CMs) stimulated by ET-1 now provides a better platform to model human-specific mechanisms of HF and test therapeutic interventions.

Alterations in L-type calcium channels (LTCCs) have been implicated in impaired excitation-contraction coupling and reduced myocardial contractility in HF. Several studies have demonstrated downregulation of LTCCs expression and reduced channel density in failing human myocardium, potentially contributing to diminished calcium influx, impaired SR calcium release, and reduced contractile function.^16,17^ However, other studies have reported no significant changes in LTCC expression or basal L-type calcium currents (I_CaL_), suggesting that alterations in channel phosphorylation, gating, and subcellular distribution may also contribute to LTCCs dysfunction in HF.^18,19^ The cardiac expression of the LTCC isoforms Cav1.2 and Cav1.3 is developmentally regulated. Although both isoforms are expressed in atrial and ventricular myocardium during fetal and neonatal development, their expression profiles diverge during postnatal maturation. Cav1.2 remains the predominant LTCC isoforms throughout the adult myocardium, whereas Cav1.3 undergoes marked developmental downregulation in the ventricles, with its expression becoming largely restricted to supraventricular tissues, including the atria, sinoatrial node, and atrioventricular node.^20^ We recently demonstrated that the C-terminus domain of Cav1.3 possess the unique ability to translocate to the nucleus and act as an auto-trans-regulator, upregulating its own gene expression, thereby restoring calcium current densities, and modulate expression of other genes.^21,22^ These findings reveal a previously unrecognized role of the Cav1.3 C-terminus in regulating cardiac calcium channel expression, which may have important implications for cardiac function and remodeling in HF. The aim of this study was to investigate the therapeutic potential of a gene-based approach to restore calcium handling and improve contractile function in a hiPSC-CMs model of HF. In this context, restoring the expression of the Cav1.3 may represent a promising therapeutic strategy to improve calcium handling and contractile function. As mentioned above, while healthy adult ventricles do not physiologically express Cav1.3, recent studies have revealed the re-expression of its mRNA and proteins in the left ventricles of patients with ischemia-induced HF, suggesting an endogenous compensatory mechanism in response of impaired contractility.^23^ collectively leveraging this dynamic by evaluating the capacity of Cav1.3-C-terminus to counteract ET-1-induced HF alterations within hiPSC-CMs, would provide preclinical validation for a novel genetic and molecular strategy to restore contractile function in HF.

## Method

### hiPSC culture, differentiation, and maintenance

One male and one female hiPSCs from Stanford University Cardiovascular Institute Biobank was used. The hiPSCs were plated in 60mm dish, coated with hESC-qualified Matrigel (Corning, NY, USA), with mTeSR^TM^ for 5 days (StemCells Technologies, Vancouver, Canada). Upon achieving a confluent growth around 80%, the cells were detached from the culture plates after a wash with PBS without calcium, followed by gentle dissociation using TrypLE^TM^ (Gibco, NY, USA) at 37°C for approximately 5 min. The hiPSCs pellet was obtained after centrifugation at 200g for 5 min, resuspend in mTeSR^TM^ with 1 µM Rock inhibitor Y-27632 (StemCells Technologies), and plated at 30,000-50,000 cells/cm^2^. Upon achieving 80% confluency, the ventricular cardiomyocyte differentiation was initiated by replacing the medium with STEMdiffTM Ventricular Cardiomyocyte Differentiation Kit (StemCells Technologies), as previously described.^24^ Cardiomyocytes derived from hiPSCs (hiPSC-CMs) exhibited spontaneous beating between day 8 and 9, after which they were maintained into STEMdiff^TM^ Cardiomyocyte Maintenance medium (StemCells technologies).

Monolayers of hiPSC-CMs were treated with 10 nM ET-1 (Bio-techne, MN, USA), for 48 hours at day 20, to induce HF-like phenotype and hypertrophic changes, as previously described.^13,14^ Another group of hiPSC-CMs were treated with ET-1, followed by infection with an AAV6 containing human Cav1.3-C-terminus (amino acids N1840 to L2137) at 5.10^3^ vg/cell on day 22 for 48 hours. The experiments were then performed at day 30.

### RT-qPCR

Total RNA was extracted from 30-day old beating hiPSC-CMs monolayers using the NucleoSpin RNA kit (Macherey-Nagel, PA, USA) and followed by a reverse transcription to obtain cDNA using the Transcriptor Universal cDNA master mix (Sigma-Aldrich, MI, USA), according to the manufacturer’s instructions. Quantitative RT-PCR was performed on a QuantStudio 5 system (Applied Biosystems, CA, USA) using the PowerUp SYBR Green Master Mix (Applied Biosystems). The target genes evaluated included atrial natriuretic peptide (*NPPA*), natriuretic peptide B (*NPPB*), skeletal α-actin (*ACTA1*), collagen type I alpha chain (*COL1A1*), potassium voltage-gated channel subfamily H member 2 (*KCNH2*), voltage-gated LTCC Cav1.3 alpha-1D subunit (*CACNA1D*), voltage-gated LTCC Cav1.2 alpha-1C subunit (*CACNA1C*), and Nav1.5 sodium channel (*SCN5A*), with the human large ribosomal protein (*RPLPO*) serving as the reference gene. The List of primers is provided in **Table 1**.

**Table 1:**
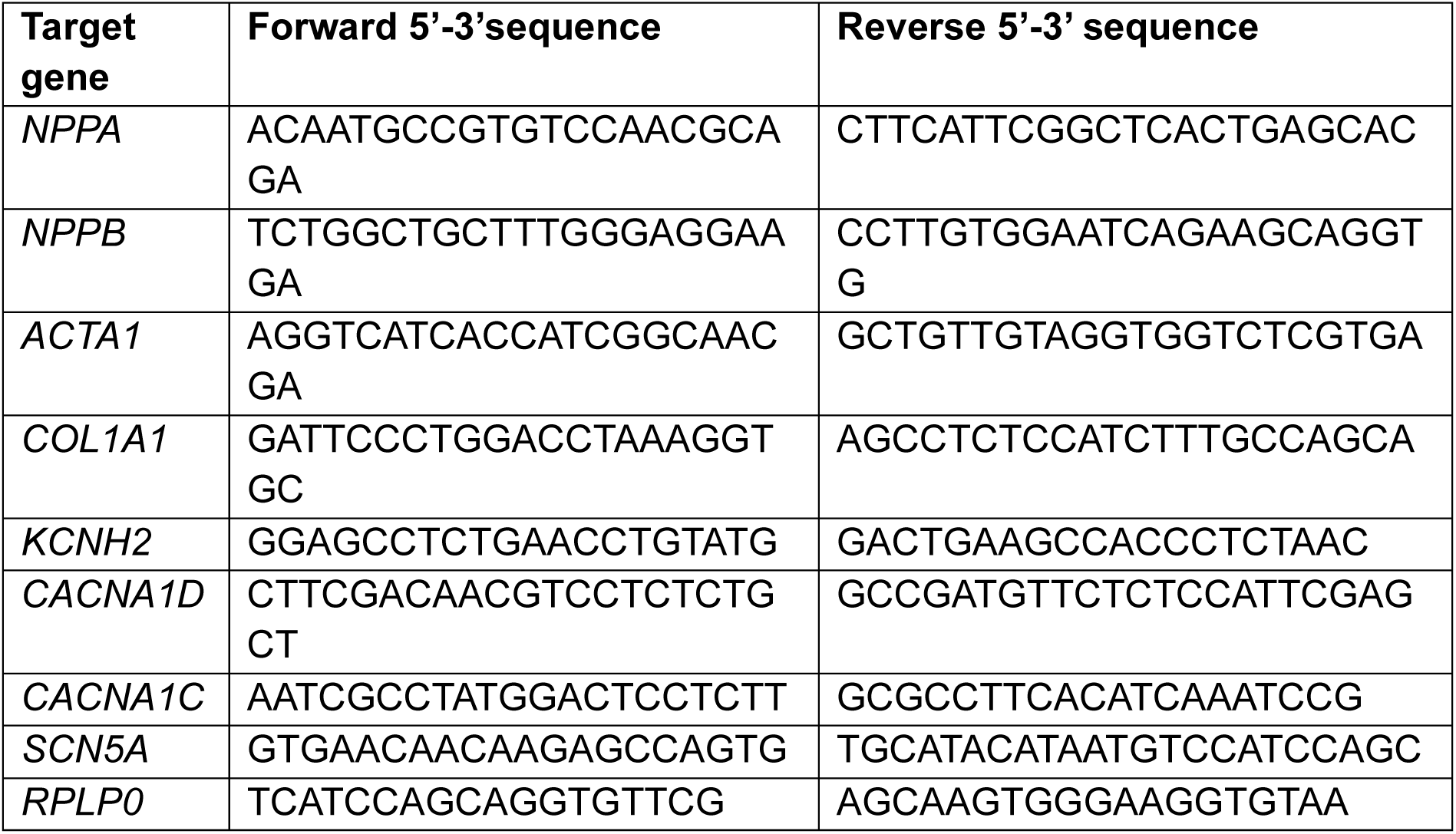
Primer sequences used in RT-qPCR.

### Electrophysiology

hiPSC-CMs were enzymatically dissociated (TrypLE^TM^) and seeded as single cells onto hESC-qualified Matrigel-coated 35 mm dishes, at least 7 days prior patch-clamp experiments. Patch-clamp experiments were performed at room temperature using Axopatch 200B amplifier (Axon Instruments, CA, USA). Pipettes were made from borosilicate glass capillaries, and fire polished.

Action potentials (APs) and potassium currents were evaluated in current clamp mode using the whole cell configuration. The patch pipettes (resistance 2-3 MOhms) were filled with a solution containing (in mmol/L): 10 NaCl, 122 KCl, 1 MgCl_2_, 1 EGTA and 10 HEPES; pH was adjusted at 7.3 with KOH. The bath solution (external solution) was composed of (in mmol/L): 154 NaCl, 5.6 KCl, 2 CaCl_2_, 1 MgCl_2_, 8 glucose and 10 HEPES; the pH was adjusted at 7.3 with NaOH. APs were elicited at 1 Hz following injection of a 3 ms, 20-1500 pA rectangular current pulse. The holding potential during the recordings was maintained at -80 mV with a Cybercyte V10 dynamic clamp system (Cytocybernetics, NY, USA). AP parameters were analyzed such as AP duration at 90%,50% and 30% of repolarization (APD_90/50/30_), the maximal upstroke velocity (dV/dtmax), and amplitude. Potassium currents were obtained using 4 sec pulses from - 40 mV to + 40mV in + 5 mV increments.

For LTCC, the patch pipettes (resistance 1-2 MOhms) were filled with (in mmol/L): 25 NaCl, 105 CsCl, 1 MgCL_2_, 10 EGTA and 10 HEPES; pH was adjusted at 7.3 with CsOH. The batch solution consisted of (in mmol/L): 100 NaCl, 5 CsCl, 5 CaCl_2_, 40 NMDG, 1 MgCl_2_, 10 glucose, 10 HEPES and 15 TEA-Cl; pH was adjusted at 7.3 with methanethiosulfonic acid. LTCC were evoked using 200 ms square steps from a holding potential of -40mV to + 35 mV at 5 mV increments.

For sodium currents, patch pipettes (resistance 1-2 MOhms) were filled (in mmol/L): 105 CsF, 35 NaCl, 10 EGTA, 10 HEPES and pH was adjusted to 7.2 with CsOH. The bath solution consisted of (in mmol/L): 105 NMDG, 35 NaCl, 2 KCl, 10 HEPES, 1 MgCl_2_, 1.5 CaCl_2_, 10 TEA-Cl and 0.5 Nimodipine with pH 7.4 adjusted with methanethiosulfonic acid. Sodium currents were obtained using 50 ms pulses from -90 mV to + 45 mV with + 5 mV increments.

APs and currents were analyzed using Clampfit (pClamp v10, Molecular devices, CA, USA).

### hiPSC-CMs contraction analysis

The contractile function of hiPSC-CMs monolayers was assessed using a custom-made video analysis software, as previously described.^25–27^ The hiPSC-CMs monolayers cultured in 12-well plates were placed on the stage of an inverted microscope (Axio Observer, Zeiss, NY, USA) equipped with Axiocam 705 (Zeiss). Phase contrast videos were acquired, with a 20x objective, at 60 frames per second. For analysis, TIFF images sequences were extracted from the videos, and the displacement of each contrasted particle was tracked frame-by-frame. This tracking generated a displacement curve as a function of time. Finally, regions exhibiting similar contractile profiles were clustered, and specific contractile parameters were quantified using a custom MATLAB script.

### Data analysis and statistics

All experiments were performed with 3 independent differentiations from each group. Data processing and statistical analyses were conducted with Prism 10 (GraphPad, MA, USA). A normality test (Agostino and Pearson omnibus normality test) was always used to determine whether data followed a normal distribution. Statistical significance was assessed using one-way ANOVA followed by Tukey’s multiple comparisons test or the Kruskal-Wallis test followed by Dunn’s multiple comparisons test, as appropriate (ns p > 0.05; *p < 0.05; **p < 0.01; ***p < 0.001).

## Results

### ET-1 induces the expression of HF-associated markers

To determine whether ET-1 treatment induces molecular changes associated with pathological cardiac remodeling, we assessed the expression of key hypertrophic and remodeling-associated genes using RT-qPCR. ET-1 treatment significantly increased the expression of the well-established cardiac hypertrophic markers *NPPA* and *NPPB* by approximately 110% each (**Fig. 1 A and B**). Similarly, the expression of *ACTA1*, another marker associated with pathological cardiomyocyte hypertrophy, was significantly increased by 144% (**Fig. 1C**), while the extracellular matrix-associated gene *COL1A1* exhibited a 166% increase (**Fig. 1D**). Interestingly, treatment with the Cav1.3 C-terminus in ET-1-treated cells showed a trend toward attenuating the upregulation of these remodeling-associated genes, although these changes did not reach statistical significance (**Fig. 1A-D**). Collectively, these findings demonstrate that ET-1 induces a molecular signature consistent with pathological cardiac hypertrophy and extracellular matrix remodeling, supporting the relevance of this model for investigating HF-associated remodeling.

**Figure 1:**
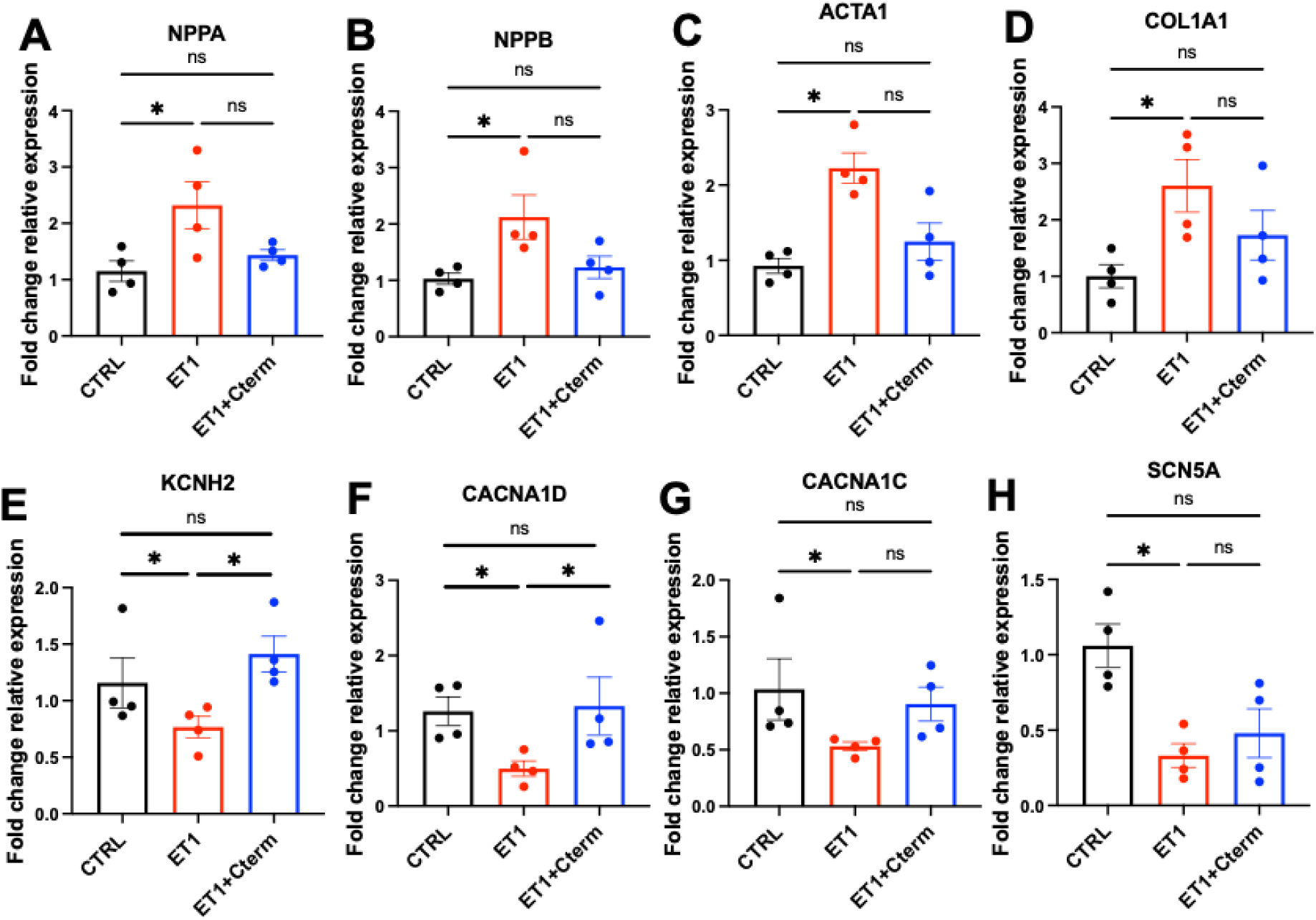
Cav1.3-C-terminus treatment partially attenuates Et-1-induced alterations in hypertrophic marker and cardiac ion channel gene expression in hiPSC-CMs. Fold change expression relative to the control group for hypertrophic markers: **(A)** atrial natriuretic peptide (*NPPA*), **(B)** natriuretic peptide B (*NPPB*), **(C)** skeletal alpha-actin (*ACTA1*), and **(D)** collagen type I alpha chain (*COL1A1).* Fold change expression relative to the control for cardiac ion channel markers: (E) Potassium voltage-gated channel subfamily H member 2 (*KCNH2*), **(F)** Cav1.3 voltage-gated L-type calcium channel (*CACNA1D*), **(G)** Cav1.2 voltage-gated L-type calcium channel (*CACNA1C*), and (H) Nav1.5 voltage-gated sodium channel (*SCN5A*). N=4 independent biological replicates; Statistical significance was assessed using one-way ANOVA followed by Tukey’s multiple comparisons test or the Kruskal-Wallis test followed by Dunn’s multiple comparisons test, as appropriate; means ± SEM; *p < 0.05. CTRL: control; ET1: endothelin-1; ET1+Cterm: Endothelin-1 + Cav1.3-C-terminus.

We next investigated whether Cav1.3-C-terminus treatment could mitigate ET-1-induced alterations in the expression of key cardiac ion channel genes associated with HF-related remodeling.

ET-1 treatment significantly downregulated *KCNH2* expression, which was restored to control levels following Cav1.3-C-terminus treatment (**Fig. 1E**). Similarly, ET-1 induced a significant reduction in the expression of both LTCCs genes, *CACNA1D* (Cav1.3) and *CACNA1C* (Cav1.2) (**Fig. 1F and G**). Notably, Cav1.3-C-terminus treatment restored *CACNA1D* expression to baseline levels, whereas *CACNA1C* exhibited a nonsignificant trend toward recovery respectively (**Fig. 1F and G**). In contrast, *SCN5A* expression was significantly reduced following ET-1 treatment and remained suppressed despite Cav1.3-C-terminus treatment (**Fig. 1H**). Collectively, these findings suggest that the Cav1.3-C-terminus treatment selectively attenuates ET-1-induced transcriptional remodeling of cardiac ion channels, particularly *KCNH2* and *CACNA1D*.

### Cav1.3-C-terminus treatment attenuates ET-1-induced electrophysiological remodeling

To assess the effects of ET-1-induced remodeling and the potential therapeutic efficacy of AAV6-mediated Cav1.3-C-terminus on the electrophysiological properties of hiPSC-CMs, we recorded APs at 1 Hz using the patch-clamp technique. Representative AP traces revealed a marked prolongation of AP duration (APD) following ET-1 treatment (**Fig. 2A**). Specifically, ET-1 significantly prolonged APD_90_ (90% of repolarization) and APD_50_ (50% of repolarization) by 34% and 17%, respectively (**Fig. 2B and C**). Notably, Cav1.3-C-terminus treatment reversed these alterations, restoring both APD_90_ and APD_50_ to control levels (**Fig. 2A-C**). In contrast, APD_30_ (30% of repolarization) remained unchanged across all experimental conditions (**Fig. 2D**). ET-1 treatment also significantly reduced the maximal AP upstroke velocity (dV/dt_max_) by 20%, an effect that was not reversed by Cav1.3-C-terminus treatment, with values remaining significantly lower than those in the control group (**Fig. 2E**). Finally, AP amplitude did not differ significantly among the experimental groups (**Fig. 2F**). Together, these findings demonstrate that Cav1.3-C-terminus treatment selectively attenuates ET-1-induced electrophysiological remodeling by restoring AP duration without significantly improving the maximal upstroke velocity.

**Figure 2:**
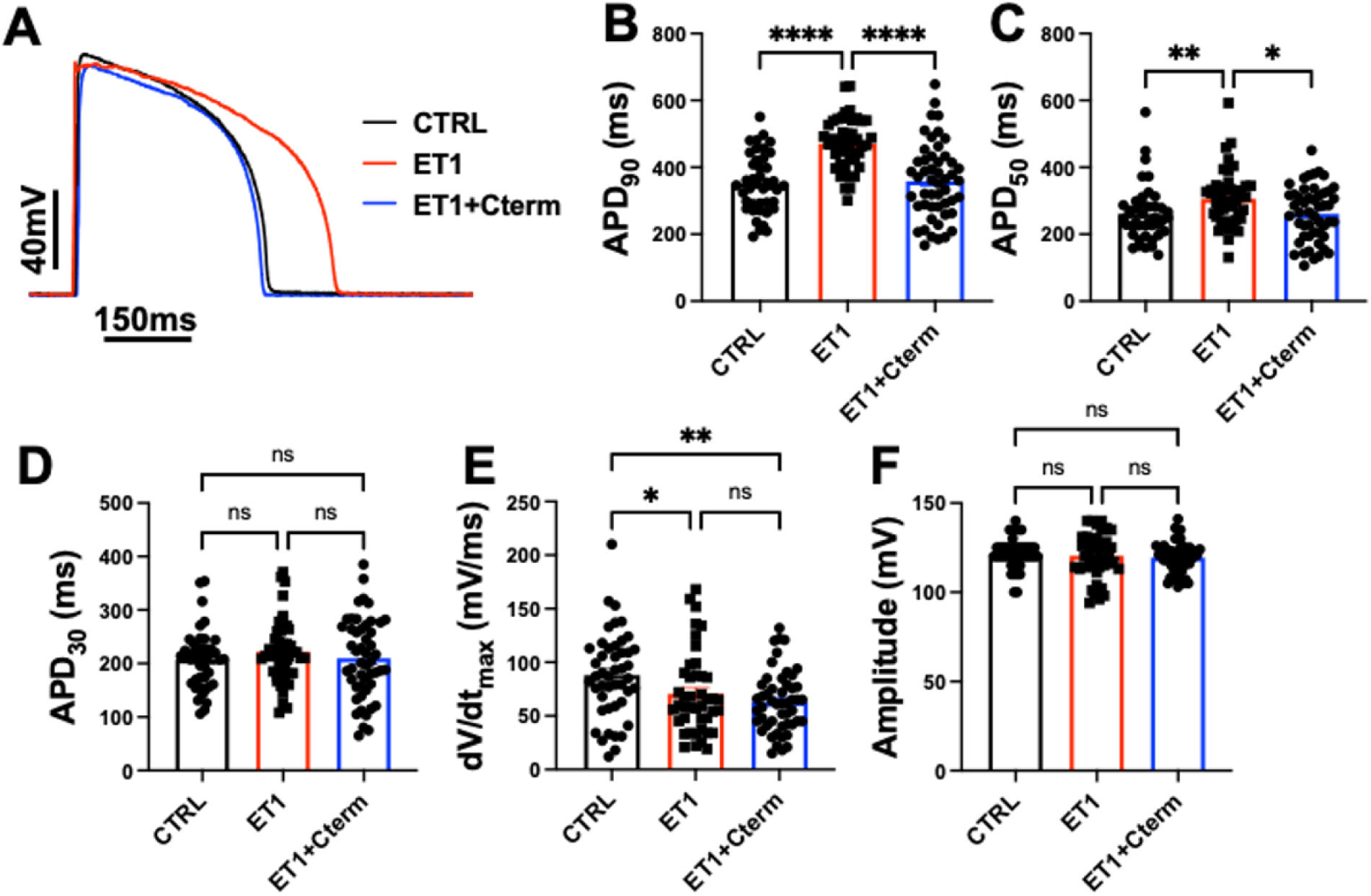
Cav1.3-C-terminus treatment reverses Et-1-induced AP prolongation in hiPSC-CMs. **(A)** Traces of action potentials (APs) elicited at 1Hz in CTRL (black; n=42), after ET-1 treatment (red; n=46), and after co-treatment ET-1 + Cav1.3-C-terminus (blue; n= 50). Measurement of AP duration at 90% of repolarization (APD_90_) **(B)**, 50% of repolarization (APD_50_) **(C)**, and 30% of repolarization (APD_30_) **(D)**. Values of maximal AP upstroke (dV/dtmax) **(E)** and amplitude of AP **(F)**. N = 3 independent differentiations for both male and female hiPSCs cell lines. Statistical significance was assessed using one-way ANOVA followed by Tukey’s multiple comparisons test or the Kruskal-Wallis test followed by Dunn’s multiple comparisons test, as appropriate; means ± SEM; *p < 0.05, **p < 0.01, **** p < 0.0001. CTRL: control; ET1: endothelin-1; ET1+Cterm: Endothelin-1 + Cav1.3-C-terminus.

We next investigated the ionic mechanisms underlying the observed alterations in AP properties. Whole-cell recordings of global outward potassium currents revealed a marked reduction in current density following ET-1 treatment, as illustrated by representative current traces (**Fig. 3A-C**). Current-voltage (I/V) relationship analysis confirmed a significant 30% decrease in outward potassium current density at +40 mV compared with control cells (**Fig. 3D**). Notably, Cav1.3-C-terminus treatment reversed this reduction, restoring potassium current density to levels comparable to those of the control group (**Fig. 3A-D**). Similarly, recordings of I_CaL_ revealed a significant 40% reduction in current density at +5 mV following ET-1 treatment (**Fig. 3H**). This decrease was reversed by Cav1.3-C-terminus treatment, which preserved calcium current density at levels comparable to those observed in control cells (**Fig. 3E-H**). Interestingly, ET-1 treatment also induced a negative shift in the voltage dependence of I_CaL_ activation, which was not reversed by Cav1.3-C-terminus treatment (**Fig. 3I**). We next assessed voltage-gated sodium currents (I_Na_). Representative I_Na_ traces (**Fig. 3J-L**) and I/V relationship analysis revealed a significant 50% reduction in I_Na_ density at -30 mV following ET-1 treatment (**Fig. 3M**). In contrast to its effects on potassium and calcium currents, Cav1.3-C-terminus treatment failed to restore I_Na_ density to control levels (**Fig. 3J-M**). Finally, to assess ET-1-induced changes in cardiomyocyte size, we measured cell membrane capacitance, an electrophysiological indicator of cell surface area and hypertrophic growth. ET-1 treatment significantly increased membrane capacitance compared with control cells (**Fig. 3N**). However, and despite its beneficial effects on AP duration and selected ionic currents, Cav1.3-C-terminus treatment did not reverse this increase, with membrane capacitance remaining significantly elevated relative to control cells and comparable to that of the ET-1-treated group (**Fig. 3N**). Collectively, these findings demonstrate that Cav1.3-C-terminus treatment selectively attenuates ET-1-induced electrophysiological remodeling by restoring potassium and L-type calcium current densities, while having no significant effect on the reduction in I_Na_ density or the increase in cell membrane capacitance. All electrophysiological results are resumed in the **Table 2**. This suggests that the beneficial electrophysiological effects of the Cav1.3-C-terminus treatment occur independently of a measurable reversal of ET-1-induced cellular hypertrophy.

**Figure 3:**
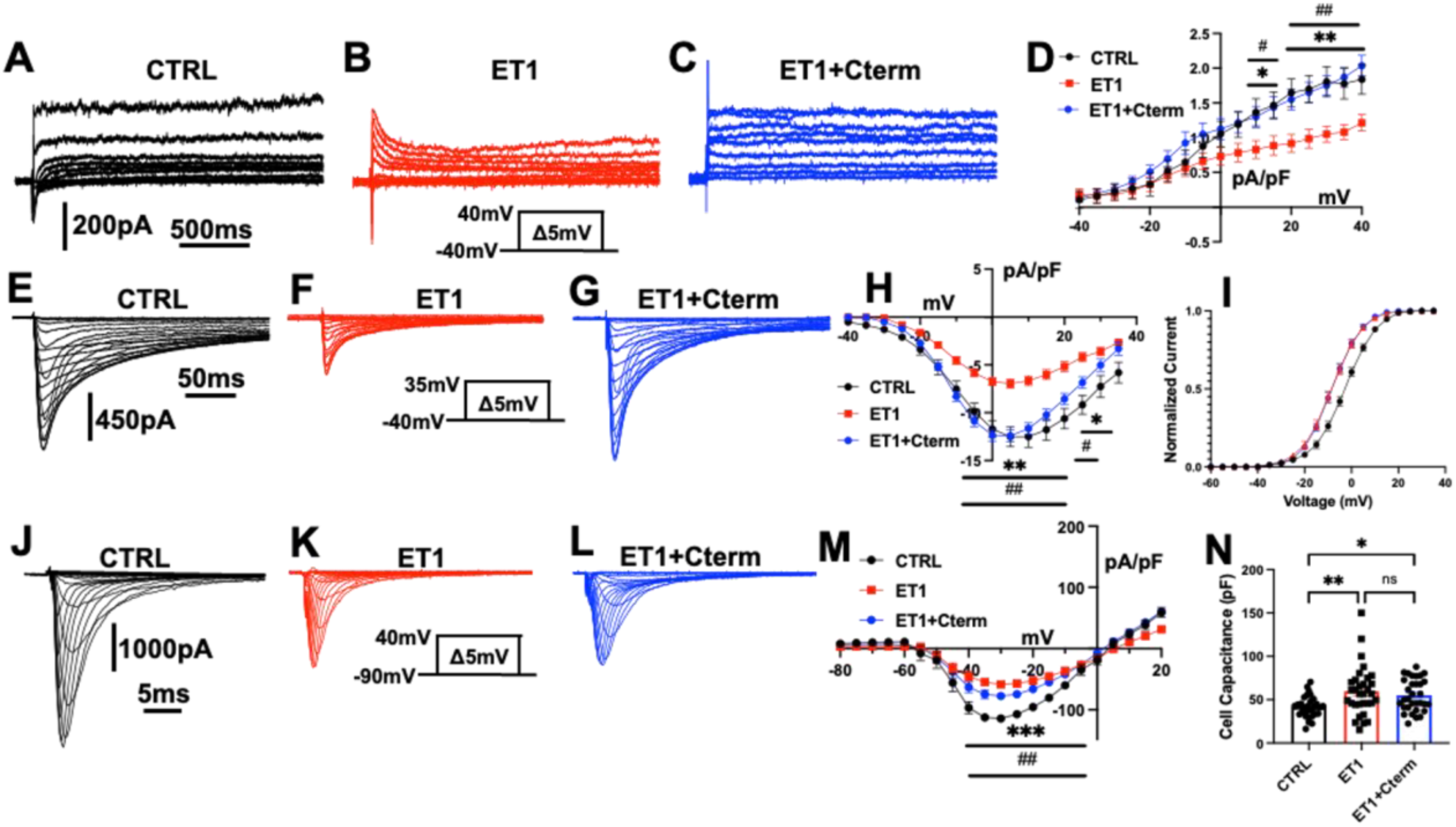
Cav1.3-C-terminus treatment restores potassium and L-type calcium currents but not sodium currents following Et-1-induced electrophysiological remodeling in hiPSC-CMs. Representative potassium currents recorded in **(A)** CTRL (black; n=19), **(B)** after ET-1 treatment (red; n=18), and **(C)** after ET-1 + Cav1.3-C-terminus co-treatment (blue; n=21), with the corresponding normalized current/voltage relationship curves (D). Representative calcium currents recorded in (E) CTRL (n=19), **(F)** after ET-1 treatment (red; n=20), and **(G)** after ET-1 + Cav1.3-C-terminus co-treatment (blue; n=22), with the corresponding normalized current/voltage relationship curves **(H)**, and steady-state activation curves **(I)**. Representative sodium currents recorded in **(J)** CTRL (n=19), **(K)** after ET-1 treatment (red; n=20), and **(L)** after ET-1 + Cav1.3-C-terminus co-treatment (blue; n=18), with the corresponding normalized current/voltage relationship curves **(M)**. Measurements of cell capacitance during voltage clamp recording **(N)**. N = 3 independent differentiations for both male and female hiPSCs cell lines. Statistical significance was assessed using one-way ANOVA followed by Tukey’s multiple comparisons test or the Kruskal-Wallis test followed by Dunn’s multiple comparisons test, as appropriate; means ± SEM; * CTRL vs ET-1; # ET-1 vs ET-1+Cterm; *^,#^p < 0.05, **^,##^p < 0.01, *** p < 0.001. CTRL: control; ET1: endothelin-1; ET1+Cterm: Endothelin-1 + Cav1.3-C-terminus.

**Table 2:** Electrophysiological results between control, endothelin-1 and endothelin-1 + Cav1.3-C-terminus treatment.

|  |  | Control | Endothelin-1 | Endothelin-1 + Cterm |
| --- | --- | --- | --- | --- |
| Action Potential |  |  |  |  |
|  | APD90 (ms) | 351 ± 14 | 470 ± 12 | 358 ± 16 |
|  | APD50 (ms) | 260 ± 13 | 305 ± 12 | 261 ± 11 |
|  | APD30 (ms) | 207 ± 8 | 222 ± 8 | 209 ± 10 |
|  | dV/dtmax (mV/ms) | 88 ± 6 | 70 ± 6 | 63 ± 4 |
|  | Amplitude (mV) | 121 ± 2 | 120 ± 2 | 119 ± 1 |
| Potassium Current |  |  |  |  |
|  | I<sub>K</sub> at +40 mV (pA/pF) | 1.8 ± 0.2 | 1.2 ± 0.1 | 2.0 ± 0.1 |
| Calcium Current |  |  |  |  |
|  | I<sub>CaL</sub> at +5 mV (pA/pF) | -12.5 ± 1 | -6.9 ± 1 | -12.4 ± 1 |
|  | V<sub>1/2</sub> (mV) | -2 ± 1 | -8 ± 1 | - 8 ± 1 |
| Sodium Current |  |  |  |  |
|  | I<sub>Na</sub> at -30 mV (pA/pF) | -114 ± 6 | -58 ± 5 | -77 ± 6 |
| Cell capacitance |  | 41.4 ± 2 | 59.7 ± 5 | 55.0 ± 3 |

### Cav1.3-C-terminus treatment restores ET-1-induced alterations in contractile function

To investigate the effects of ET-1-induced remodeling and the potential therapeutic benefits of Cav1.3-C-terminus treatment on the functional properties of hiPSC-CMs, we assessed the contractile behavior of spontaneously beating hiPSC-CM monolayers. Representative displacement traces illustrate the contraction profiles of control, ET-1-treated, and ET-1 with Cav1.3-C-terminus-treated monolayers (**Fig. 4A-C**). Quantitative analysis revealed that ET-1 treatment significantly reduced the spontaneous beating rate by 44% compared with the control group (**Fig. 4D**). Notably, Cav1.3-C-terminus treatment completely reversed this chronotropic effect, restoring the beating rate to control levels (**Fig. 4D**). Similarly, ET-1 treatment markedly impaired contractile function, as evidenced by a significant 70% reduction in contraction amplitude (**Fig. 4E**). Consistent with the chronotropic effect, Cav1.3-C-terminus prevented the ionotropic property, maintaining the contraction amplitude at levels comparable to those observed in control group (**Fig. 4E**). In contrast, neither contraction time (**Fig 4F**) nor relaxation duration (**Fig. 4G**) differed significantly among the experimental groups. All contractile properties measurements are resumed in **Table 3**. Collectively, these findings demonstrate that Cav1.3-C-terminus treatment effectively attenuates ET-1-induced contractile dysfunction, restoring spontaneous beating rate and contraction amplitude without significantly altering contraction or relaxation kinetics.

**Figure 4:**
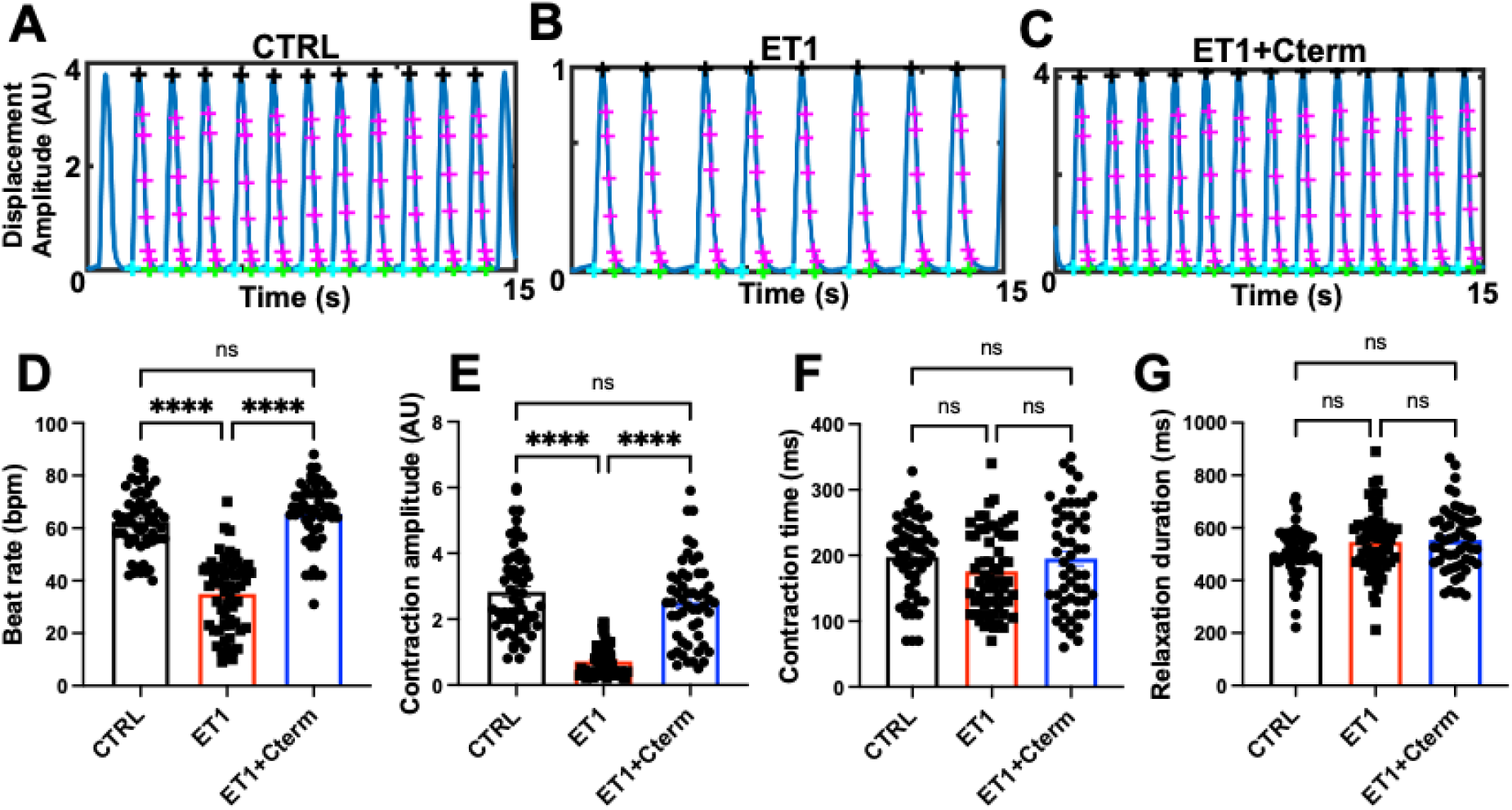
Cav1.3-C-terminus treatment restores spontaneous beating rate and contraction amplitude following Et-1-induced contractile dysfunction in hiPSC-CMs. Representative contraction cycles from phase-contrast videos obtained in hiPSC-CMs monolayers in CTRL (n=58) (A), after ET-1 treatment (n=61) (B), and after co-treatment with ET-1 and Cav1.3-C-terminus (n=53) (C). Corresponding contractions parameters measurement: spontaneous beating rate (D), contraction amplitude (E) and time (F), and relaxation duration (G). N = 3 independent differentiations for both male and female hiPSCs cell lines. Statistical significance was assessed using one-way ANOVA followed by Tukey’s multiple comparisons test or the Kruskal-Wallis test followed by Dunn’s multiple comparisons test, as appropriate; means ± SEM; **** p < 0.0001. CTRL: control; ET1: endothelin-1; ET1+Cterm: Endothelin-1 + Cav1.3-C-terminus.

**Table 3:** Contractile properties between control, endothelin-1 and endothelin-1 + Cav1.3-C-terminus treatment.

|  | Control | Endothelin-1 | Endothelin-1 + Cterm |
| --- | --- | --- | --- |
| Beat Rate (bpm) | 62 ± 2 | 35 ± 1 | 65 ± 2 |
| Contraction Amplitude (A.U.) | 2.8 ± 0.2 | 0.7 ± 0.1 | 2.5 ± 0.2 |
| Contraction Time (ms) | 197 ± 8 | 175 ± 8 | 195 ± 10 |
| Relaxation Duration (ms) | 502 ± 12 | 546 ± 15 | 553 ± 17 |

## Discussion

The present study demonstrates that ET-1 treatment effectively recapitulates key HF hallmarks in hiPSC-CMs, including molecular, electrophysiological, and contractile alterations, supporting their utility as a human-relevant *in vitro* model of pathological cardiac remodeling. hiPSC-CMs represent a promising alternative approach to animal models, providing a platform for investigating disease mechanisms, identifying potential biomarkers, and evaluating novel therapeutic strategies. Building on this model, we investigated the therapeutic potential of Cav1.3-C-terminus treatment in mitigating HF-associated electrophysiological and contractile dysfunction. Notably, Cav1.3-C-terminus treatment restored the expression of *CACNA1D*, which encodes the Cav1.3 L-type calcium channel, and *KCNH2*, accompanied by improvements in ionic currents, AP duration, and contractile function. Collectively, these findings highlight the therapeutic potential of Cav1.3-C-terminus treatment in attenuating pathological electrophysiological remodeling and restoring key aspects of cardiomyocyte function in an *in vitro* model recapitulating HF hallmarks.

The classic hallmarks of HF, and particularly HFrEF, include structural and functional features such as cardiomyocytes hypertrophy, fibrosis, and decreased contractility.^28,29^ Interestingly, our ET-1-based HF model effectively recapitulates these features. ET-1 treatment increased the gene expression of hypertrophic markers such as *NPPA, NPPB*, and *ACTA1*, which corresponded with an increase in cell capacitance measured during patch-clamp experiments. This confirms a previous study showing that ET-1 induces cardiac hypertrophy in neonatal rat cardiomyocytes.^30^ While earlier studies, using ET-1 to mimic HF in hiPSC-CMs, focused solely on contractile function, the present study explores the subsequent cellular electrophysiological modifications in greater detail.^10,14^ We found that ET-1 also drives electrophysiological remodeling by decreasing both calcium and potassium currents, which aligns with existing literature.^31^ However, in contrast to a previous study demonstrating a stimulatory effect of ET-1 on sodium currents, our results showed a decrease in sodium current amplitude.^32^ This discrepancy can likely be explained by the acute ET-1 application used in their research. Ultimately, the reduction of these key ion currents prolongs the APD, which increases arrhythmia susceptibility by forming functional conduction blocks or triggering early afterdepolarizations (EADs).^33,34^

ET-1 contribute to disturb both chronotropic and inotropic effects of hiPSC-CMs monolayer, like previously observed in a HF model.^13^ The spontaneous activity of hiPSC-CMs is fundamentally calcium-dependent, disturbance of calcium entry in hiPSC-CMs contribute to this decrease of chronotropic parameters observed after ET-1 treatment.^35,36^ For the ionotropic response, HF induce a downregulation of LTCC expression and function in ventricular myocytes.^37^ This reduction contributes to smaller calcium transients and contractile weakness in HF. Furthermore, combined with the severe structural remodeling, mark a disruption and loss of T-tubules, which physically distances the remaining LTCC from RyR2.^38,39^ This critical uncoupling delays and desynchronizes calcium release, drastically reducing contractile efficiency and serving as a primary driver of systolic dysfunction in failing hearts.

This is the primary rationale for utilizing the Cav1.3-C-terminus to drive the re-expression of the Cav1.3 LTCC and restore normal calcium handling in the failing heart. Although treatment with the Cav1.3-C-terminus failed to reverse cardiomyocyte hypertrophy in hiPSC-CMs, its restorative effects on electrophysiological properties and contractile function are highly significant. We previously demonstrated that in an ischemia-induced murine HF model, the Cav1.3-C-terminus upregulated Cav1.3 gene expression and increased LTCC density, which contributed to restoring the ejection fraction and preventing arrhythmias.^22^ These findings are corroborated by the present study in an HF-like hiPSC-CMs model, where the application of the Cav1.3-C-terminus successfully normalized calcium current density and restored normal chronotropic and inotropic parameters. Interestingly, the present study demonstrates that the Cav1.3-C-terminus restores not only the gene expression and current densities of LTCCs, but also those of potassium channels. These results raise the question of whether the Cav1.3-C-terminus acts directly on potassium channel gene expression, specifically *KCNH2* in this context, or if this is an indirect effect. Both mechanisms are plausible. A previous study demonstrated a direct upregulation of calcium-activated potassium (SK2) channels by the Cav1.3-C-terminus, suggesting a similar direct mechanism could apply to *KCNH2*.^40^ On the other hand, an indirect upregulation is also conceivable due to the strong correlation between *CACNA1C* and *KCNH2* expression in adult human heart tissue and hiPSC-CMs, where increase of *CACNA1C* expression affect the gene expression of *KCNH2.*^41^ This correlated expression is crucial for reducing the incidence of proarrhythmic events. Furthermore, because calcium acts as a second messenger for signaling networks like the calcium-calmodulin pathway, the normalization of calcium handling itself could indirectly restore potassium gene expression.^42^

The use of hiPSC-CMs provides substantial advantages but also entails certain limitations. A major challenge in hiPSC-CM models is the lack of cellular morphologic and electrophysiologic maturity, which is accompanied by fetal-like molecular features.^43^ To address this issue, we employed a dynamic clamp technique to maintain the resting potential at levels comparable to those of adult human cardiomyocytes. Depending on the specific research objectives, the heterogeneous composition of hiPSC-CMs, comprising ventricular-, atrial-, and nodal-like cardiomyocyte subtypes, as well as the limited cellular complexity of cardiomyocyte monocultures, may present experimental challenges and influence the physiological relevance of the model. Nevertheless, hiPSC-CMs remain a valuable platform in cardiovascular research, with broad applications ranging from elucidating the genetic and molecular mechanisms underlying cardiac diseases to investigating electrophysiological alterations and evaluating pharmacological responses.

In summary, the present study, together with previous findings, demonstrates the feasibility of recapitulating key HF hallmarks in hiPSC-CMs through ET-1-induced pathological remodeling. ET-1 application promoted the upregulation of cardiac hypertrophic markers and induced pronounced electrophysiological alterations and contractile dysfunction, reproducing several characteristic features of HF. Importantly, this model provides a valuable platform for evaluating novel therapeutic approaches, as demonstrated by AAV6-mediated delivery of the Cav1.3-C-terminus. Notably, Cav1.3-C-terminus treatment attenuated ET-1-induced electrophysiological remodeling by restoring *CACNA1D* and *KCNH2* expression, normalizing calcium and potassium current densities, and reversing AP prolongation. These electrophysiological improvements were accompanied by the restoration of spontaneous beating rate and contraction amplitude in hiPSC-CM monolayers, indicating recovery of key chronotropic and inotropic parameters. Collectively, these findings identify Cav1.3-C-terminus treatment as a promising gene-based therapeutic strategy for mitigating HF-associated electrophysiological and contractile dysfunction, warranting further investigation in more physiologically complex models of HF.

## Data availability

The data supporting this article and other findings are available within the manuscript, figures, supplementary figures and from the corresponding author upon request.

## Acknowledgements

We thank the platform Neurophotonics from Québec, Canada that provided the AAV6 and plasmid containing the Cav1.3-C-terminus construct. CTRL hiPSCs cell line was obtained as a generous gift from the Stanford Cardiovascular Institute (SCVI) Biobank.

We would like to acknowledge our fundings from Biomedical Laboratory Research & Development Service of United States (U.S.) Department of Veterans Affairs, Office of Research and Development, Merit Review grant I01 BX002137 to MB; National Heart, Lung, and Blood Institute 1R01HL164415 to MB; US Department of Defense award number W81XWH-21-1-0424 to MB. This work was also supported by an American Heart Association Postdoctoral Fellowship award number 25POST1361676 to J-BR.

## Author contributions

Y.S and J-B.R. designed the experiments, contributed to the discussion, and wrote the manuscript. J-B.R. and Y.S collected and analyzed the data. Y.S., J-B.R. and M.B., contributed to the discussion and reviewed/edited the manuscript.

